# Bayesian population receptive field modeling in human somatosensory cortex

**DOI:** 10.1101/577981

**Authors:** Alexander M. Puckett, Saskia Bollmann, Keerat Junday, Markus Barth, Ross Cunnington

## Abstract

Somatosensation is fundamental to our ability to sense our body and interact with the world. Our body is continuously sampling the environment using a variety of receptors tuned to different features, and this information is routed up to primary somatosensory cortex. Strikingly, the spatial organization of the peripheral receptors in the body are well maintained, with the resulting representation of the body in the brain being referred to as the somatosensory homunculus. Recent years have seen considerable advancements in the field of high-resolution fMRI, which have enabled an increasingly detailed examination of the organization and properties of this homunculus. Here we combined advanced imaging techniques at ultra-high field (7T) with a recently developed Bayesian population receptive field (pRF) modeling framework to examine pRF properties in primary somatosensory cortex. In each subject, vibrotactile stimulation of the fingertips (i.e., the peripheral mechanoreceptors) modulated the fMRI response along the post-central gyrus and these signals were used to estimate pRFs. We found the pRF center location estimates to be in accord with previous work as well as evidence of other properties in line with the underlying neurobiology. Specifically, as expected from the known properties of cortical magnification, we find a larger representation of the index finger compared to the other stimulated digits (middle, index, little). We also show evidence that the little finger is marked by the largest pRF sizes. The ability to estimate somatosensory pRFs in humans provides an unprecedented opportunity to examine the neural mechanisms underlying somatosensation and is critical for studying how the brain, body, and environment interact to inform perception and action.

## 1. Introduction

Mechanoreceptors permeate the human body and serve as key communicators between the body and the brain. They are ubiquitous near the very boundary of the body, embedded throughout the skin (Horch et al., 1977; Vallbo and Hagbarth, 1968). They are also distributed deep within the body, being found in articular tissues such as joint capsules and menisci (Zimny, 1988; Zimny et al., 1988). As such, mechanoreceptors are responsible for responding to information about both the external environment (i.e., exteroception) and about the state of the body itself (i.e., proprioception). The signals from these peripheral receptors are transmitted via the spinal cord to somatosensory cortex; the processing there being fundamental to our sensation of touch (Kandel et al. 2000). Information, originating from the various receptors, is then further fed forward to be utilized by a greater network of cortical areas (Mauguiere et al., 1997). This network of areas integrates the somatosensory information with other sensory and motor information critical for haptic perception as well as a wide range of sensorimotor tasks necessary for interacting with the environment (Haegens et al., 2011; Lederman and Klatzky, 2009).

A great deal of scientific work has been done to understand the organization and function at each stage of processing between the mechanoreceptors and the cortex. For this, recordings have been made in the periphery, directly from single nerve fibers carrying information from cutaneous receptors (Johansson, 1978), as well as from various stages in the central nervous system (Celesia, 1979; Ibanez et al., 1992). These studies have spanned animal (Fleetwood-Walker et al., 1988; Liu et al., 2013) and human models (Vallbo and Johansson, 1984) and have drawn upon a wide variety of both invasive (Jeanmonod et al., 1989) and non-invasive (Davis et al., 1998) measurement techniques. In humans, it has been shown that the signals from mechanoreceptors are routed through the dorsal horn and the thalamus, where some lower-order processing occurs, before reaching the cortex for higher-order processing. One striking aspect of the organization of this system is that the spatial relationship among the receptors in the body is conserved along this journey between the body and the brain (Hong et al., 2011; Yamada et al., 2007), the consequence of which is the presence of an orderly, somatotopically organized representation of the body in primary somatosensory cortex – i.e., the sensory homunculus (Schott, 1993).

The modern-day concept of the sensory homunculus originated from the neurological work of Wilder Penfield and Edwin Boldrey (Penfield and Boldrey, 1937). Published in 1937, Penfield and Boldrey presented summary data from the electrical stimulation of sensorimotor cortex in 126 surgical patients – finding an orderly map of the body within the brain. They depicted this using a distorted drawing of the human body, with the distortions reflecting the amount of cortex associated with the somatosensory or motor functions of the depicted body part. This concept was named the homunculus (Latin for “little man”), and has significantly impacted scientific research in the field and related neurosurgical practice since (Catani, 2017). Although many aspects of the homunculus are still under debate (e.g., degree of specificity / overlap among neighboring somatotopic locations, boundary between motor and somatosensory areas, and individual variability in somatotopic maps), what is clear is that the basic spatial organization of the receptors in the body is reflected in the cortex.

Not only is the spatial organization of the mechanoreceptors represented in an orderly fashion within the brain, but the amount of cortex dedicated to each body part has been shown to generally correspond to the density of innervation – and perhaps more importantly – the behavioral relevance of that body part (Catania and Henry, 2006). Moreover, the response characteristics of cortical neurons in somatosensory cortex are similar to the mechanoreceptors in the periphery. Pertinently, as is the case with the mechanoreceptors of the body (Johansson, 1978), these neurons do not respond to a single location in body space, but are instead, characterized by a topographic sensitivity profile – i.e., a receptive field (RF).

Although measuring RF properties from peripheral nerves is possible in healthy human volunteers as it is minimally invasive, measuring somatosensory RF properties within the cortex has been mainly restricted to animal models and patient populations (e.g., those already planned to undergo surgery (Lenz et al., 1988)). Consequently, it has been difficult to examine and compare the response properties throughout each stage of somatosensory processing in awake and behaving humans. This has begun to shift, however, with the invention and subsequent refinement of non-invasive neuroimaging techniques. Basic demonstrations of tactile stimulation eliciting cortical activation within human S1 were shown using Positron Emission Tomography (PET) (Fox et al., 1987; Greenberg et al., 1981). Using functional magnetic resonance imaging (fMRI), it later became possible to resolve this activity with such detail that the responses could be attributed to the stimulation of individual fingers (Francis et al., 2000; Gelnar et al., 1998). More recently, high-resolution fMRI has borne evidence that human S1 actually contains multiple orderly somatotopic maps of the fingers, both across (Martuzzi et al., 2014; Sanchez-Panchuelo et al., 2010) and within (Sanchez-Panchuelo et al., 2012) digits.

It is evident that high-resolution fMRI is closing the gap between the electrophysiological-based recordings and non-invasive estimates of cortical RF properties. Being able to use fMRI to map the organization of S1, for example, shows its ability to estimate the somatotopic location of each imaging voxel’s receptive field. Other measures such as a voxel’s response profile to stimulation of body space on and around the center of its receptive field (Besle et al., 2014; Martuzzi et al., 2014) can been seen as estimates of the size of that voxel’s RF. It is important to note here that a voxel’s RF is more properly referred to as its population receptive field (pRF). This distinction is critical as the pRF of a voxel is the estimate of the receptive field properties of a summed population of neurons (i.e., all the neurons within the volume of an imaging voxel), rather than the RF of a single neuron. With this knowledge and thoughtful experimental designs, however, it is possible to non-invasively gain unprecedented insight into the receptive field properties of the neurons contained within each voxel.

Here we extend this line of research by using previously collected, high-resolution fMRI somatotopic mapping data (Puckett et al., 2017) with a novel Bayesian pRF modeling framework (Zeidman et al., 2018) to demonstrate the feasibility of using vibrotactile driven sensory responses in S1 to directly estimate each voxel’s pRF. The pRF modeling approach marks an improvement over conventional phase-encoded techniques (Puckett et al., 2017; Sanchez-Panchuelo et al., 2010) by providing an estimate of not only the preferred fingertip (pRF center location) but also the size and shape (i.e., the topography) of the pRF. Moreover, the Bayesian approach to pRF modeling has advantages over the traditional pRF technique (Dumoulin and Wandell, 2008) by providing estimates of the uncertainty associated with the pRF parameter estimates, by accounting for variability in the hemodynamic response across the brain, and by providing a formal framework to test competing pRF models (e.g., Gaussian vs. Difference of Gaussian or symmetrical vs. asymmetrical profiles).

## 2. Materials and Methods

### 2.1 Subjects

Six, right-handed subjects (23-31 years, mean 27 years) with no history of neurological or psychiatric diseases completed the original experiment (Puckett et al., 2017). The experiment was conducted with the written consent of each subject and was approved by the local ethics committee in accordance with national guidelines.

### 2.2 Stimulation and tasks

Here we used data from only one of the experimental conditions (i.e., the sensory condition) from the original study to perform the pRF mapping. During this condition, tactile stimulation was delivered via a MR-compatible, piezoelectric, vibrotactile stimulator (www.hybridmojo.com). The device consisted of 4 units, each able to deliver vibrotactile stimulation to the pad (i.e. volar surface) of a single fingertip. The stimulation timing and frequency could be controlled independently for each unit.

During each run, the 4 fingertips (index, middle, ring, and little) of the right hand were sequentially stimulated using a phase-encoded design (Besle et al., 2013; DeYoe et al., 1996; Engel, 2012; Engel et al., 1994; Sereno et al., 1995). For this, each individual fingertip was stimulated for 7872 ms before moving to the next. Each cycle of stimulation began with the index finger and ended with the little finger. Stimulation then returned to the index finger to begin another stimulation cycle. The frequency of stimulation changed every 1968 ms (synced with the MRI scanner repetition time), and three frequencies were used (5, 20, and 100 Hz). The stimulation frequency was programmed to change randomly among the three frequencies, except that the same frequency could not occur twice in a row at a fingertip. Each run was comprised of 5 cycles of stimulation (31.5 s in duration each).

### 2.3 Magnetic resonance imaging data acquisition

Data were acquired on a MAGNETOM 7T whole-body research scanner (Siemens Healthcare, Erlangen, Germany) with a 32-channel head coil (Nova Medical, Wilmington, US). Whole-brain, anatomical images were collected using an MP2RAGE sequence (Marques et al., 2010) with a TE of 2.88 ms, TR of 4300 ms, flip angles of 5 and 6 degrees, TI_1_ of 840 ms, TI_2_ of 2370 ms, FOV of 201 mm x 224 mm x 144 mm, and a matrix size of 378 × 420 × 288 -resulting in an isotropic voxel size of 0.5 mm.

Functional data were collected using a 3D-EPI sequence (Poser et al., 2010) with a blipped CAIPIRINHA (Breuer et al., 2006; Setsompop et al., 2012) implementation (Poser et al. 2014a; Poser et al. 2014b; Zahneisen et al., 2015). Scan parameters were as follows: TE of 30 ms, TR of 82 ms, flip angle of 17 degrees, echo spacing of 0.97 ms, FOV of 160 mm x 160 mm x 39 mm, and a matrix size of 192 × 192 × 48 – resulting in an isotropic voxel size of 0.8 mm. The acquisition was accelerated by a factor of 2 in-plane and by a factor of 2 in the slice-encoding direction with a CAIPI-shift of 1 using the GRAPPA (Griswold et al., 2002) image reconstruction pipeline as provided by the vendor – resulting in a total acceleration factor of 4 and an effective volume TR of 1968 ms. The acquisition slab was positioned obliquely to ensure adequate coverage of S1 in the left hemisphere, contralateral to the stimulated fingertips. 12 runs of stimulation were collected in a single scan session yielding approximately 1 hour of data per subject to be used for the pRF modeling. Periods of baseline fMRI activity were also measured during each run with the sensory condition beginning and ending with a 31.5 s block of rest (no tactile stimulation). The 5 cycles of stimulation occurred between these blocks.

### 2.4 Preprocessing

MRI data were pre-processed using the AFNI/SUMA analysis package (Cox, 1996; Saad and Reynolds, 2012) as follows: volume registration of the functional data, alignment of the anatomical and the functional data, averaging of time courses, removal of baseline periods, and then smoothing. For volume registration, each EPI volume was registered to the minimum outlier fraction volume (i.e. the volume that is least different from all the others after detrending). To bring the anatomical and functional data into alignment, the anatomical dataset was skull-stripped and then aligned to this same EPI base using AFNI’s align_epi_anat.py script. The time-courses for all 12 runs of stimulation were then averaged at each voxel across the repetitions, and the baseline periods were removed. To increase signal-to-noise while maintaining the spatial resolution necessary to resolve cortical representations of individual fingertips (Martuzzi et al., 2014), the images were smoothed using a 1.2 mm Gaussian kernel.

### 2.5 Previous delay analysis

For details on the original delay analysis see our previous publication (Puckett et al., 2017). Because we compare the results from the Bayesian pRF modeling approach to that from the delay analysis, a brief summary of this analysis is provided here. The fMRI response delay was calculated at each voxel using a Phase estimator based on the Hilbert transform (Saad et al., 2003) as implemented in AFNI’s Hilbert Delay plugin. For each voxel, this analysis returns the correlation coefficient (cc) and response delay at which the correlation between the empirical time-course and the reference waveform is maximum. The reference waveform was a sine wave with five cycles and a 31.5 s period matching the timing of the movement of sensory stimulation, which is swept across all four fingertips five times (i.e. for five cycles) with each cycle being 31.5 s in duration.

### 2.6 Bayesian pRF modeling

#### 2.6.1 Overview

The pRF modeling was performed using the BayespRF Toolbox (available from https://github.com/pzeidman/BayespRF), which is dependent on Matlab (here we used version R2018b) and SPM (here we used version 12, available from http://www.fil.ion.ucl.ac.uk/spm). The BayespRF Toolbox was designed to provide a generic framework for mapping pRFs associated with stimulus spaces of any dimension onto the brain, but it was only evaluated by the developers for mapping 2-dimensional (2D) visual pRFs in human visual cortex (Zeidman et al., 2018). Here we modified and applied the toolbox to examine mapping somatosensory pRFs in human S1.

We adhered to the basic procedures outlined in the original publication associated with the BayespRF Toolbox (Zeidman et al., 2018), utilizing the following two scripts supplied with the toolbox: Run_first_level.m and Run_pRF_analysis.m. The first level of analysis (Run_first_level.m) prepares the data for the pRF modeling procedure, mainly by reducing the number of voxel time-courses to model and hence the time required for the modeling computations. Within Run_first_level.m, this is achieved by performing a general linear model (GLM) analysis in SPM. Only data from voxels surviving threshold are then taken forward for the actual pRF modeling (per Run_pRF_analysis.m).

#### 2.6.2 Modifications for somatosensory space

In order for the procedures to be suitable for our somatosensory data, some modifications were required at both stages of the original analysis (i.e., GLM and pRF modeling). The major modification required at the GLM stage was simply that of re-defining the task regressors. For the original visual pRF analysis, Run_first_level.m was set up with 9 task-related regressors. These were defined by dividing the visual field into 9 equal squares, and then building regressors based on the timing of visual stimulation within those 9 subfields. Here, we modified this by defining only 4 regressors – one per fingertip.

At the pRF modeling stage, there were two main modifications required of the original analysis: (1) that of defining the stimulus space and (2) that of constraining the pRF parameters. In the original analysis, the stimulus space was defined in terms of degrees of visual angle and the limits were matched to the stimulus display. Here we defined the somatosensory space using the same 2D matrix but with arbitrary dimensions limited to ±10 in both dimensions and divided along the x-axis into 4 segments of equal width (representing each individual fingertip). It is important to note that the data we have can only be used to map across 1 dimension in this 2D sensory space. The nature of our stimulators is such that the entire volar surface of each fingertip is stimulated before moving to the next digit, and hence, our data can only be used to map the across-digit dimension. However, within-digit somatotopy has, with the use of more spatially specific stimulation, been shown to run perpendicular to the across-digit dimension within the cortex (Sanchez-Panchuelo et al., 2012). For this reason, we kept the 2D representation of somatosensory space and addressed the limited nature of our data by constraining the pRF centers in one of the two dimensions (at y = 0). Along with the use of symmetrical pRF models, this reduces the 2D problem to 1D (i.e., we only estimate location and size in the across-digit dimension). We did not place any constraints on the center location in the across-digit dimension (i.e., the center could be continuously distributed anywhere between x = ±10). Constraints were also placed on the pRF size with the minimum size not being allowed to be less than 1/10^th^ of the sensory space occupied by a single fingertip, and the maximum size restricted to the equivalence of all four fingers (i.e., 20 units). While it is possible that some of the modeled voxels have pRFs that extend beyond the four fingertip representations in somatosensory space, we would not be able to resolve these given our experimental design.

#### 2.6.3 Application and voxel selection

As mentioned, the first level of analysis was a simple GLM designed to reduce the number of voxel responses to be modeled by removing those voxels without task-related signals. Only data from voxels surviving threshold (*p < 0.05*, uncorrected) were taken forward for pRF modeling. The threshold at this first level was set liberally in order to prevent the exclusion of weak or potentially unusual signals that might still be able to be successfully modeled – at the cost of increased compute time. Surviving voxels were then submitted to the second level of analysis, i.e., the pRF modeling. The main goal of this step was to optimize, on a voxel-wise basis, the fit between an estimated waveform and the empirically measured BOLD time-course by modifying the position and size of the pRF model. Following the procedure of Zeidman etl al. (2018), a second threshold was applied after the pRF modeling at a posterior model probability > 0.95. Voxels surviving this threshold were used for data visualization. Finally, data were restricted to only include voxels in primary somatosensory cortex. For this, we used the same S1 ROI as in our previous publication (defined using the independent, phase-delay analysis) (Puckett et al., 2017). Together, this resulted in the final set of voxels contributing to our pRF estimates.

### 2.7 Surface reconstruction and data visualization

Cortical reconstruction and volumetric segmentation were performed using FreeSurfer, which is freely available for download (http://surfer.nmr.mgh.harvard.edu/) (Dale et al., 1999; Dale and Sereno, 1993). Data were projected onto a computationally-inflated surface model using AFNI/SUMA. To map the data from volume to surface domains the volumetric data were sampled at 10 evenly spaced points between the white matter and pial surfaces. The most common value along each segment (i.e., the mode) was mapped onto the corresponding node of the inflated surface model. Note that the cortical surface models were only used for data visualization and region-of-interest (ROI) definition. All analyses and statistics were performed using the volumetric data.

## 3. Results

### 3.1 Overview

Vibrotactile stimulation of the fingertips elicited a patch of BOLD activation in primary somatosensory cortex, along the post-central gyrus, in all subjects. We previously analysed these signals using a phase-delay technique revealing somatotopic organization with individual fingertip specificity within this patch (Fig. 1A) (Puckett et al., 2017). Here, we reanalyzed these signals using a recently established Bayesian pRF modeling framework (i.e., the BayespRF Toolbox) to investigate the possibility of estimating somatosensory pRFs from high-resolution fMRI data. We found, that with only minor modifications, the BayespRF Toolbox could be used to successfully model pRFs in S1. Examining the estimated pRF centers (Fig. 1b, top) reveals a nearly identical somatotopic map as that produced with the phase-delay approach. Whereas the delay analysis only provides estimates of each voxel’s preferred fingertip (effectively its pRF center), the Bayesian modeling approach also provides estimates of the pRF size (Fig. 1B) as well as a number of neuronal and hemodynamic parameter (Fig. 1C).

**Figure 1.**
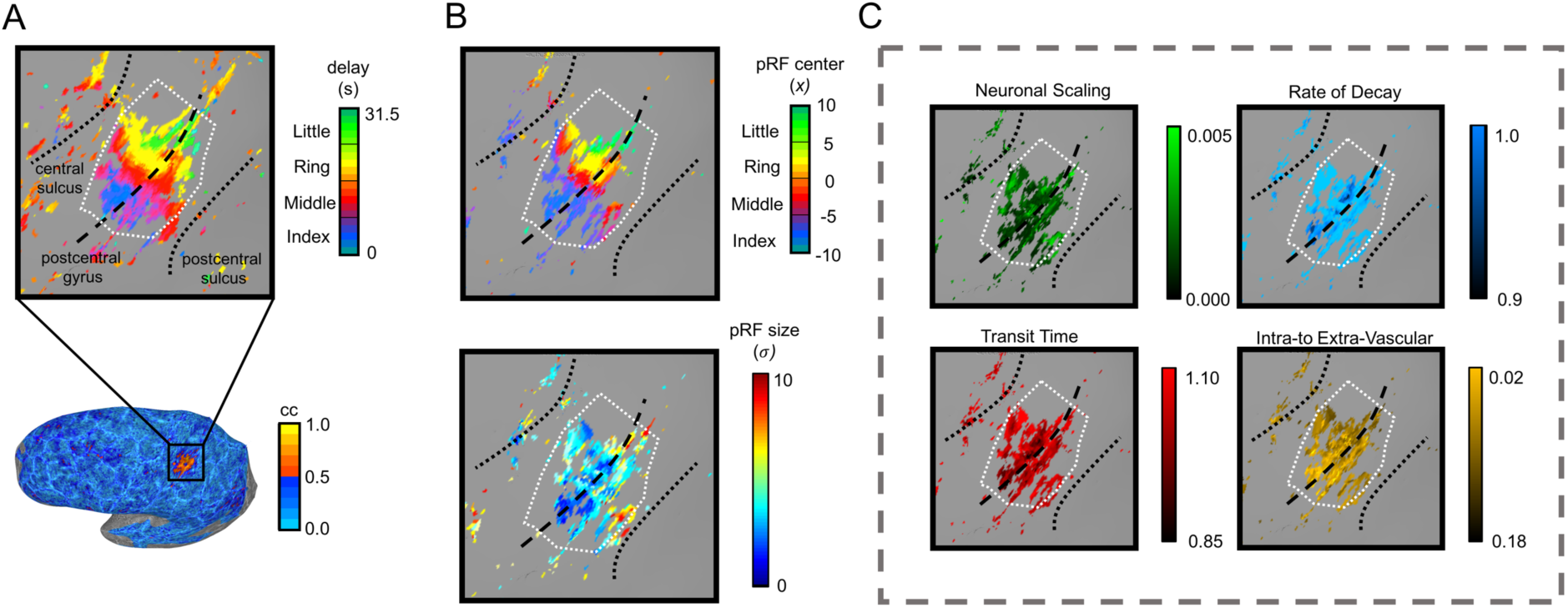
Activation in primary somatosensory cortex resulting from vibrotactile stimulation of individual fingertips. (A) Results from the previous phase-delay analysis showing the presence of an across-digit, somatotopic map in S1 (top), and the associated correlation map (bottom) for anatomical orientation (adapted from Puckett et al., 2017). (B) Results from the Bayesian modeling analysis. Color represents the pRF center location in the top map and pRF size in the bottom. (C) In addition to pRF parameters, the Bayesian approach also provides voxel-wise estimates of a number of neuronal and hemodynamic parameters, shown here projected onto the cortical surface model (scaling of neuronal response, transit time, rate of decay, and ratio of intra-to extra-vascular signal). Note that this data is from Subject 1, the pRF was modeled using a Gaussian response profile, and the white dashed line represented the S1 ROI boundary.

### 3.2 Bayesian pRF analysis

As described in section 2.6, the pRF modeling analysis consisted of two levels (GLM and pRF modeling stages) followed by the application of an S1 ROI to select the final set of voxels used to examine the pRF estimates (voxel counts at the various stages of analysis can be found supplementary Table S1). Figure 2 illustrates the single voxel modeling results for three different voxels. For each, there is a depiction of the prior and the posterior pRF models along with their respective predictive density (PD) distributions, which represent the uncertainty in the pRF position and width. For example, the prior PD was computed by averaging the responses across 1000 samples taken from the model’s prior multivariate distribution over the parameter space. The prior PD associated with each voxel in Figure 2 is characterized by a distribution stretching across the x-dimension (across-digit) but centered and focused at y=0. This reflects the fact that we constrained the pRF parameter space to be appropriate given our stimulation, which was only applied in the across-digit dimension (see section 2.6 for details). Importantly, the large degree of across-digit uncertainty visible in the prior PD for each voxel (Fig. 2) has been greatly reduced after the modeling procedure (evident in the more punctate distribution of the posterior PDs).

**Figure 2.**
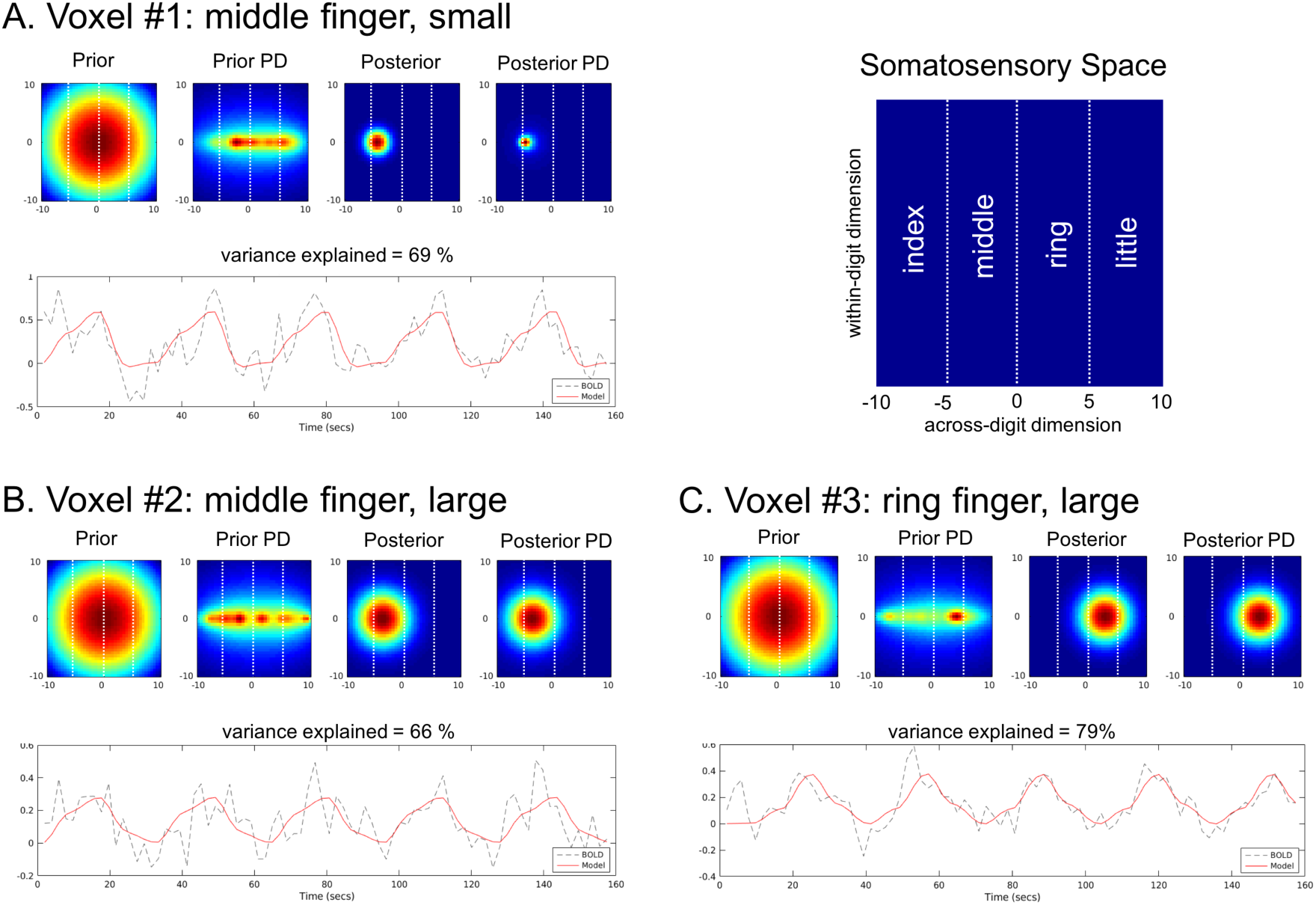
Modeling results for three S1 voxels (A, B, and C) as well as a schematic of the representation of somatosensory space (upper right). For each voxel, the prior and the posterior pRF models are shown on top, along with their respective predictive density (PD) distributions which represents the degree of uncertainty in the pRF models. Vertical dashed white lines denote the separate digit representations. Below the pRF plots is the modeled waveform (red solid line) atop the empirical BOLD time-course (black dashed line). Note that the variance in the empirical time-course explained by the model is also shown. Data is from Subject 1.

The close fit between the model and the data suggested by the reduction in uncertainty between the prior and posterior PDs can also be seen by inspection of the single voxel time-courses. Below the pRF estimates in Figure 2 are two traces showing the modeled waveform (red, solid line) atop the empirical BOLD time-course (black, dashed line) for that voxel, along with the percent variance explained by the modeled waveform. Note the close correspondence between the two traces as well as the high-degree of variance explained. For further interpretation, see the schematic of somatosensory space in the upper right of Figure 2 and recall that the pRF center could be distributed anywhere between x = ±10 and each fingertip was defined as occupying an equal amount of that space (i.e., 5 units along the x-axis with fingertips ordered from index-middle-ring-little). Together then, inspection of the estimated pRFs shows that the first two voxels (Fig. 2A and B) have pRFs with similar center locations (middle finger) to one another but different sizes, whereas the third voxel’s pRF (Fig. 2C) has a similar size to the second but a different center location (ring finger).

In addition to the pRF parameters, the modeling procedure also estimates various neuronal and hemodynamic parameters (Fig. 1C). Zeidman et al., (2018) showed a practical benefit in allowing these parameters to vary on a voxel-by-voxel basis over the use of a canonical model (nearly 20% of voxels showed strong evidence in favor of the model with free parameters). This approach has a strong theoretical foundation as well given that it has been shown that hemodynamic response varies significantly across many factors such as subjects (Aguirre et al., 1998; Handwerker et al., 2004), days (Neumann et al., 2003), age (Jacobs et al., 2008), and brain region (Birn et al., 2001, Puckett et al., 2014). Although there is clear variability in hemodynamic and neuronal parameters present in our data, we had no explicit hypotheses regarding this variability. As such, the data presented in Figure 1C are primarily for illustrative purposes – with further analyses restricted to the pRF parameters only (i.e., center location and size).

### 3.3 Somatosensory pRF parameters

In agreement with our previous analysis, the pRF center estimates show an orderly representation of the fingertips along the post-central gyrus in response to vibrotactile stimulation in all subjects (Fig. 1B and Fig. 3). It can be seen that pRF center maps and the phase-delay maps from the previous analysis produce very similar somatotopic maps (cf. Fig. 1A and Fig. 1B for a single subject example; cf. Fig. 3 here and Fig. 3 in (Puckett et al., 2017) for all subjects). In addition to the pRF center location, the Bayesian modeling approach also provides estimates of the pRF sizes (Fig. 1B and Fig. 3). Qualitatively, the cortical surface maps of pRF size appear similar among all subjects, with the exception of Subject 4, which appears to contain a higher proportion of large pRFs relative to the other subjects. Interestingly, in the other subjects there appears to be a banding pattern that runs parallel to the digit representations suggesting that the pRF sizes might vary in a digit specific manner.

**Figure 3.**
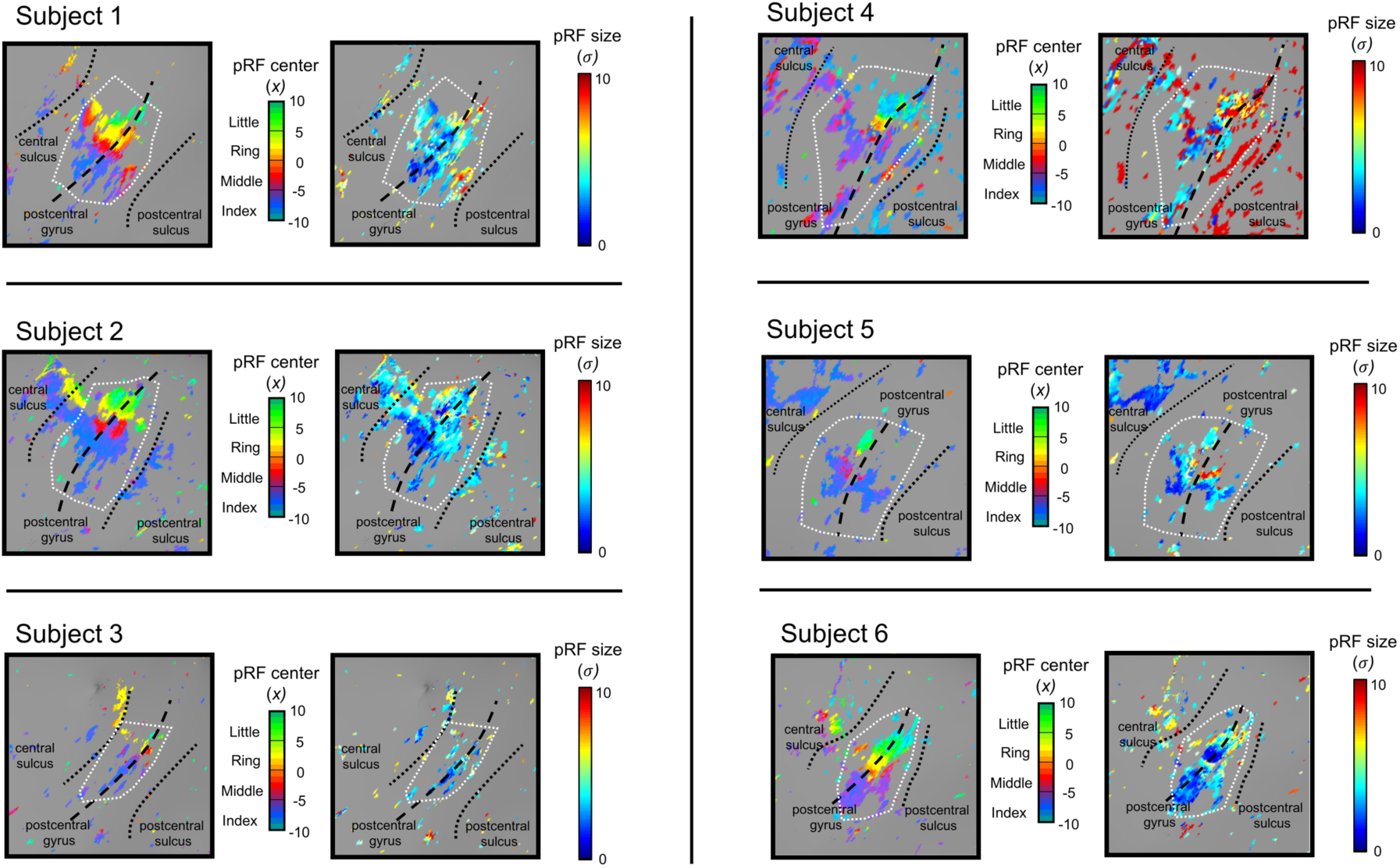
Cortical surface maps of the pRF parameters. For each individual subject, the pRF center locations (left) and the pRF sizes (right) are shown in S1 (zoomed in on the post-central gyrus, see Fig. 1A for anatomical orientation). White dashed line illustrates the ROI boundary.

To more quantitatively assess the pRF parameters, histograms were constructed at the individual and group level (Fig. 4, light grey). At the individual level, the histograms were made from voxel counts with the pRF centers binned according to each of the four digits and the pRF size binned per unit of somatosensory space. At the group level, histograms of pRF parameters were also constructed but represented in terms of the probability density rather than raw voxel counts. Inspection of the histograms reveals that variability exists at the individual subject level, yet it does appear that certain features seen within individual subjects emerge at the group level as well. Of particular note is the disproportionate number of voxels dedicated to the index finger compared to the others (middle, ring, little). The pRF size estimates tend to be distributed between x = 0 and 10 and skewed toward the smaller sizes in that range. However, a small population of voxels appear to have pRF size estimates distributed between x = 15 and 20. To interpret these pRF size estimates, recall that each finger is defined as occupying 5 units of the somatosensory space, and hence, the entire somatosensory space being modeled here for the four fingertips spans 20 units.

**Figure 4.**
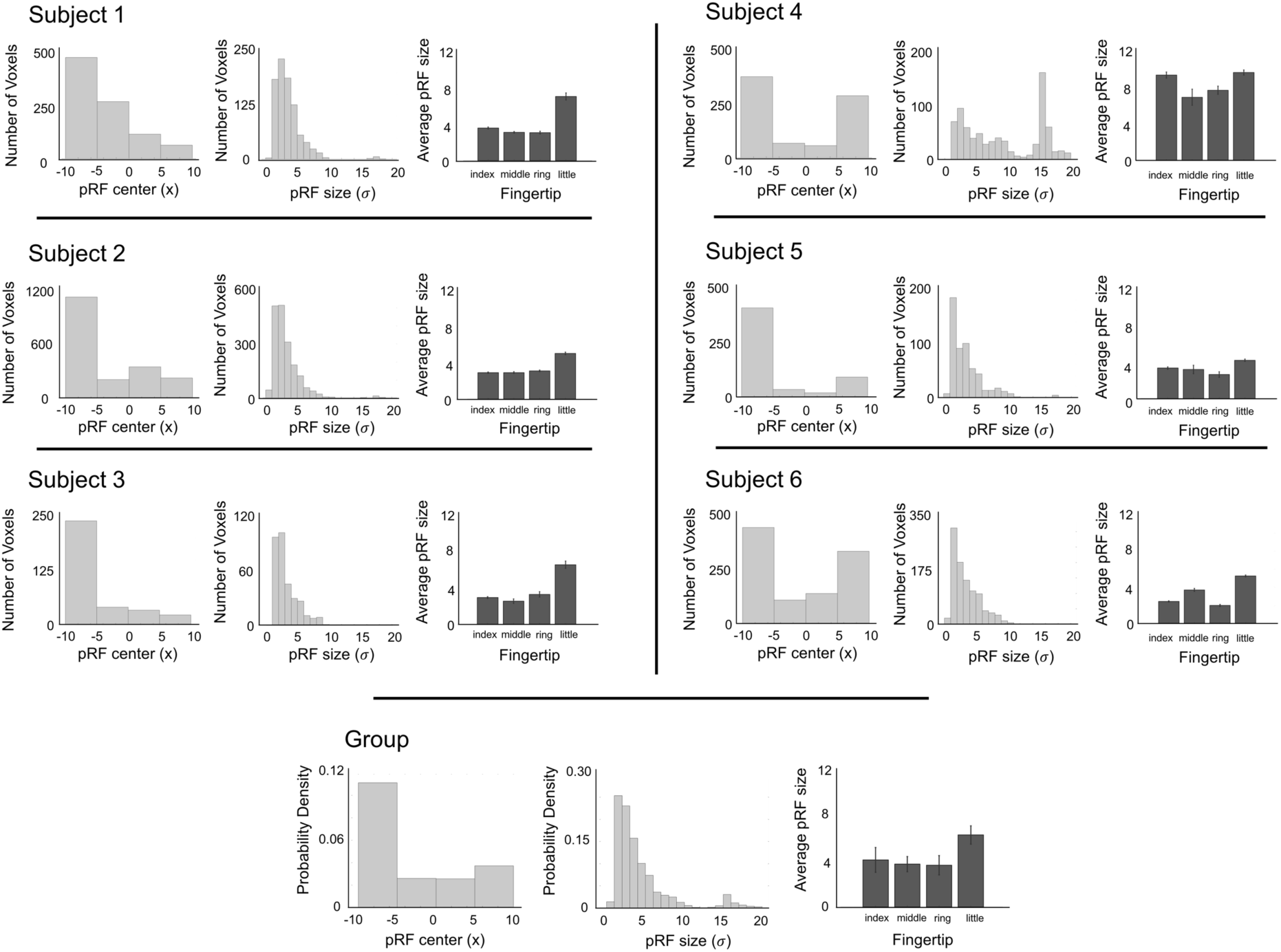
pRF parameters at the individual subject and group levels. Histograms of pRF center location and size are illustrated as the light grey graphs. Average pRF size per binned fingertip are illustrated as the dark grey graphs – error bars represent SEM across voxels at the individual level and across individuals at the group level.

In addition to the histograms, we computed the average pRF size per fingertip. This was done at the individual level from all surviving S1 voxels and at the group level by taking the mean of the average pRF size per fingertip across the individual subjects. Inspection of these graphs (Fig. 4, dark grey) for the individual subjects suggests that the pRF size does, in fact, vary according to digit, and this is supported by finding that the pRF centers and sizes were significantly correlated across voxels within 5 of 6 individuals (*p < 0.001* for Subjects 1, 2, 3, 5, and 6; *p = 0.78* for Subject 4). At the group level, the most salient characteristic of this relationship evident in Figure 3 is that the little finger appears to be marked by larger pRFs than the other three digits.

### 3.4 Gaussian vs. Difference of Gaussians pRF model

A number of visual pRF mapping studies (including that by Zeidman et al. using the Bayesian approach) have shown that some voxels in visual cortex are better modeled with a Difference of Gaussians (DoG) function compared to a single Gaussian model (Zeidman et al., 2018; Zuiderbaan et al., 2012). The main difference being that the DoG effectively incorporates a suppressive zone around the Gaussian’s excitatory center. Because the DoG model has additional parameters compared to the Gaussian model (i.e., is fundamentally more complex), testing for the most appropriate model type often involves applying some sort of information criterion after the pRF analysis (e.g., Akaike’s information criterion) (Akaike, 1974; Puckett and DeYoe, 2015); however, one of the strengths of the Bayesian pRF modeling approach is that the estimation procedure directly provides an approximation of the model evidence – the negative variational free energy (*F*). The free energy term increases with model accuracy and decreases with model complexity, and can hence be used to compare pRF models in order to determine the most accurate, least complex explanation of the data.

To assess whether the DoG function might also better model the pRFs in somatosensory cortex, we reran the entire pRF modeling analysis but with a symmetrical DoG pRF profile. Afterwards, we inspected the pRF center maps produced using a DoG pRF model, finding – that as expected – both the Gaussian and DoG models produced nearly identical maps (see Supplementary Fig. S1 for an example). Next, to determine which model best accounted for the data, we compared the *F* values at the individual and group levels. For individual subjects, we performed *t*-tests between the free energy values for all the voxels that survived threshold for both the Gaussian and DoG analyses (see supplementary Table S1 to see the proportion of these joint voxels). In doing so, we found a higher *F* value associated with the Gaussian model for all 6 subjects with this difference being statistically significant in 5 of these 6 subjects (*p ≤ 0.005* for Subjects 1, 2, 3, 5, and 6; *p = 0.12* for Subject 4) – in favor of the Gaussian model. However, this did not survive at the group level when comparing the average *F* values for each subject; there was no statistical difference at the group level between the two model types (*p = 0.13*).

## 4. Discussion

### 4.1 Overview

This study used high-resolution fMRI at 7T and a recently established Bayesian framework (i.e., the BayespRF Toolbox) to estimate pRFs in somatosensory cortex. Vibrotactile stimulation of the fingertips drove BOLD response modulation in S1, along the post-central gyrus. These responses were then used to estimate the size, location, and topography of the pRFs in S1. We were able to successfully model pRFs associated with all four of the stimulated fingertips, in all subjects. We found more voxels with pRF center locations at the index finger than the other three digits (middle, ring, little). We also found that pRF size correlated with the center location – with the little finger marked by larger pRFs than the other digits. Evidence was found within individual subjects suggesting that the pRFs in somatosensory cortex estimated using our stimulation paradigm are better characterized by a simple, excitatory Gaussian profile than one that incorporates a suppressive surround (i.e., a DoG profile), although this was not confirmed by a statistical test at the group level.

### 4.2 The somatosensory population receptive field

Somatosensory cortex is responsible for processing information from a number of different sensory receptors distributed throughout the body including mechanoreceptors, thermoreceptors, nociceptors, and chemoreceptors (Kandel et al. 2000)f. Given the nature of our stimulation (i.e., vibrotactile), we expect the responses measured in S1 to primarily be driven by activation of cutaneous mechanoreceptors. However, there are multiple types of mechanoreceptors, each with different receptive field properties. There are four main types of mechanoreceptors in the glabrous skin of humans: Merkel disc receptors, Meissner (or tactile) corpuscles, Pacinian (or Lamellar) corpuscles, and Ruffini (or Bulbous) corpuscles. Of these, the Merkel, Meissner, and Pacinian receptor types all respond to different frequencies of mechanical stimulation whereas the Ruffini corpuscles are primarily responsive to skin stretch related to mechanical deformation within joints (Grigg and Hoffman, 1982). The slowly adapting Merkel cells are most sensitive to low frequency stimulation (10 Hz), whereas rapidly adapting Meissner corpuscles are most sensitive to vibrotactile frequencies of 30 Hz, and Pacinian corpuscles are most sensitive to high-frequency vibrations around 200 Hz (Friedman et al., 2004). Given that our stimulation continuously changes across a wide range of frequencies (5, 20, and 100 Hz), we expect that our pRF measurements reflect a mixture of all three of these receptor types.

The receptive field properties of the peripheral receptors have been well characterized (Johansson, 1978; Vallbo and Johansson, 1984). For example, we know that both Merkel and Meissner receptors have smaller pRFs than the Pacinian receptors. However, because the pRFs we estimate likely result from the stimulation of a mixture of different receptor types it is difficult to validate the results by comparing them directly to the known receptive field properties of specific peripheral receptors. Nonetheless, a number of our findings are in agreement with the known organization and response properties of the somatosensory system. The estimated pRF centers are in agreement with our previous analysis (Puckett et al., 2017) as well as other published work (Maldjian et al., 1999; Martuzzi et al., 2014; Sanchez-Panchuelo et al., 2010) showing a mediolateral ordering of digits along the post-central gyrus. In line with the known properties of cortical magnification (Duncan and Boynton, 2007; Sutherling et al., 1992), our results also show a disproportionate number of voxels with pRFs centers associated with the index finger compared to the other digits.

We are aware of only one other published study that has reported pRF estimations in somatosensory cortex measured using fMRI (Schellekens et al., 2018). There are, however, two crucial differences between that study and the one here. First, the experiment by Schellekens et al. was designed to investigate pRF properties in motor cortex, not somatosensory. As such, the cortical responses were not driven by applied sensory stimulation but instead by movement of the digits. Under these conditions the authors were able to estimate pRFs in M1 (although these may better be referred to as “response” fields rather than “receptive” fields). In addition, they found an orderly map of pRFs in S1, presumably driven by the activation of deeper, proprioceptive receptors which respond to movement of finger joints rather than the more superficial mechanoreceptors targeted here (Edin, 1990). The second significant difference between this study and ours is methodological with Schellekens et al. using the conventional pRF approach rather than the Bayesian approach employed here. Despite these differences, we see similar results across the two studies. Specifically, we report the same spatial distribution of pRF center locations as well as larger pRF sizes for the little finger compared to the other three digits.

### 4.3 Behavioral relevance

The three different types of mechanoreceptors contributing to our pRF estimates are known to be linked with different aspects of tactile perceptions (i.e. pressure, flutter, and vibration). The slowly adapting Merkel cells have been linked to perceptions of pressure, texture, and the form of an object, rapidly adapting Meissner corpuscles appear to be integral to the perception of flutter, slip, and motion of objects, and Pacinian corpuscles are most sensitive to the perception of vibration (Friedman et al., 2004). Moreover, the tactile thresholds associated with each receptor type, and hence associated perceptive abilities, are known to vary (Ferrington et al., 1977). Being able to directly estimate somatosensory pRFs will provide opportunity to examine the relationship between pRF properties and these various tactile perceptions.

It is important to understand that pRF properties are not only relevant to the processing of different forms of bottom-up, sensory driven information, but that they also influence top-down effects such as attention. Findings have shown that attention modulates the responses of neurons with tactile receptive fields centered on an attended stimulus (Hsiao et al., 1993), and we have previously shown using high-resolution fMRI that the attentional field (AF) is able to modulate somatotopically appropriate regions of cortex with a fine level of detail (i.e., with individual fingertip specificity) (Puckett et al., 2017). In fact, the authors of a recent review on somatosensory attention suggested that one key advantage of having a detailed neural representation of the body in the brain is so that attention can leverage the topographical organization to select stimuli based on their somatotopic location (Gomez-Ramirez et al., 2016). The exact nature of the somatosensory attentional field and how it interacts with pRFs, however, remains an active and important area of research. A larger amount of work investigating the interaction between RFs and AFs has been performed in visual cortex compared to somatosensory cortex, where it has been shown that the relative sizes of the RF, AF, and visual stimulus appear to influence what type of attentional modulation occurs (e.g., contrast-gain vs. response-gain) (Reynolds and Heeger, 2009). Empirical measurements of somatosensory pRFs will hence provide important data that can be used to test for similar effects in somatosensory cortex, ultimately, contributing to an understanding of the neurophysiological basis of the perceptual effects associated with somatosensory attention.

### 4.4 Limitations and future directions

This work clearly demonstrates the feasibility of using vibrotactile stimulation of peripheral mechanoreceptors to map pRFs in somatosensory cortex, but it is not without limitations. Addressing these limitations can help direct further development, and as such, we discuss a few of the potential future directions here. Perhaps the greatest limitation of the current study is the spatially coarse nature of the applied sensory stimulation. The vibrotactile stimulators used here are only capable of delivering stimulation to the entire volar surface of each individual fingertip. This effectively limits the ability to resolve very small pRFs as any receptive field smaller than an individual digit would be fully activated when stimulating that digit. The solution here is only a matter of engineering a MR-compatible device capable of administering more spatially specific stimulation, and work is already being done in this direction. For example, Dancer Design (http://www.dancerdesign.co.uk/) currently builds an MR-compatible device capable of delivering vibrotactile stimulation to an area of ∼1mm^2^. Using such a device would not only permit the fingers to be stimulated at a finer spatial scale in the across-digit dimension, but it would also permit stimulating multiple sites along each finger (i.e., mapping the within-digit dimension). In fact, a previous study did just this using the Dancer Design stimulator finding an orderly representation of the within-digit dimension running orthogonal to the across-digit dimension (Sanchez-Panchuelo et al., 2012). Positioning these small stimulators across both across-and within-digit dimensions would thus permit the pRFs to be more completely characterized (e.g., by allowing one to test if the pRFs are symmetrical in both dimensions).

Using stimulation that would permit mapping across both across-and within-digit dimensions would also permit pRFs to be compared across the sub-regions of S1 as the within-digit mapping permits accurate delineation of these sub-regions (Sanchez-Panchuelo et al., 2012). The S1 ROI used here almost certainly contains multiple somatosensory areas, corresponding to the four cytoarchitectonically defined areas: 3a, 3b, 1 and 2 (Brodmann 1909; Vogt and Vogt 1919). It has traditionally been held that these areas are tailored for specific functions and are differentially sensitive to the stimulation of different receptors (e.g. deep vs. cutaneous) (Iwamura et al., 1993; Powell and Mountcastle, 1959). They are also hierarchically organized with pRF size and feature complexity increasing as one progresses up this hierarchy (Bodegard et al., 2001; Iwamura, 1998). Being able to non-invasively measure the response properties within these sub-regions brings with it the opportunity to quantitatively examine their differences and subsequently relate them to human perception and behavior.

We see several potential applications of this technique; for example, one of the more obvious extensions of this line of research would be to examine pRFs encoding somatosensory space other than the four fingertip representations (i.e., the thumb, the face, the body, etc.). fMRI is already being used to map these other locations (Sanchez Panchuelo et al., 2018), and these endeavors would undoubtedly benefit from the richer data provided by the pRF approach compared to the more typical, phase-encoded or event-related approaches. Another particularly interesting extension of this work would be to examine the feasibility of mapping pRFs from specific mechanoreceptor types. As mentioned, there exist four main types of mechanoreceptors in human skin and these have been shown to have different receptive field profiles when measuring from peripheral nerves. Although these differences are relatively minor between some receptor types, they are substantially different for others. For example, Pacinian corpuscles have RFs with only one zone of maximal sensitivity and the sensitivity profile changes gradually across the RF (similar to a Gaussian profile). However, the Meissner corpuscles and Merkel receptors are characterized by having multiple zones of maximal sensitivity and the sensitivity diminishes quickly with increasing distance away from these zones (Johansson, 1978). As mentioned above, our vibrotactile stimulation likely drives activity in all three of these receptor types. But by using specific frequencies of vibrotactile stimulation it may be possible to bias the pRFs toward certain mechanoreceptor classes. Similarly, it is reasonable to expect that this technique could be used to estimate pRFs associated with somatosensory receptors other than mechanoreceptors. For example, it has been shown that detailed maps of the digits can be measured in S1 using fMRI when applying nociceptive-selective laser stimuli to the hand (Mancini et al., 2012). Combining this type of stimulation with a pRF mapping procedure should enable the nociceptive-related pRFs to be estimated. Finally, laminar differences in somatosensory RFs have been reported from invasive measurements in the macaque (Sur et al., 1985), and applying the pRF modeling procedure to sub-millimeter data suitable for laminar fMRI (Huber et al., 2018; Lawrence et al., 2017; Puckett et al., 2016) may permit investigation of cortical-depth dependent pRF differences in humans.

## 5. Conclusion

We show that it is possible to non-invasively estimate pRFs in primary somatosensory cortex using high-resolution fMRI at 7T and a freely available Bayesian pRF modeling toolbox. This was accomplished by passing vibrotactile stimulation across the individual fingertips to activate peripheral mechanoreceptors and corresponding neuronal populations in somatosensory cortex. The ability to estimate somatosensory pRFs in humans provides an exceptional opportunity to examine the cortical representation of the body in the brain, the response properties therein – and ultimately the cortical processes underlying somatosensation.

## Acknowledgments

We thank Aiman Al-Najjar, Nicole Atcheson, and Steffen Bollmann for help with data collection, and the authors acknowledge the facilities of the National Imaging Facility (NIF) at the Centre for Advanced Imaging, University of Queensland. This work was supported by the Australian Research Council (DE180100433) and the National Health and Medical Research Council (APP 1088419). M.B. acknowledges funding from Australian Research Council Future Fellowship grant FT140100865, and S.B. acknowledges support through the Australian Government Research Training Program Scholarship.

## Supplementary Material

**Table S1.** Voxel counts after the GLM, pRF modeling, and ROI restriction for all subjects. The raw datasets contained 1,769,472 voxels. The term “joint” refers to common voxels between the Gaussian and DoG analyses, within the S1 ROIs.

| Subject | Voxel Count |  |  |  |  |  |
| --- | --- | --- | --- | --- | --- | --- |
|  | After<br>GLM | After<br>pRF Modeling |  | Within<br>ROI |  |  |
|  |  | Gaussian | DoG | Gaussian | DoG | Joint |
| 1 | 73,758 | 12,798 | 12,342 | 881 | 914 | 752 |
| 2 | 37,873 | 11,657 | 12,168 | 1,834 | 1,834 | 1,751 |
| 3 | 25,536 | 4,113 | 3,807 | 333 | 323 | 289 |
| 4 | 30,290 | 8,449 | 10,823 | 769 | 788 | 680 |
| 5 | 30,325 | 4,793 | 4,997 | 541 | 516 | 488 |
| 6 | 30,978 | 6,600 | 6,284 | 1,010 | 943 | 905 |

**Figure S1.**
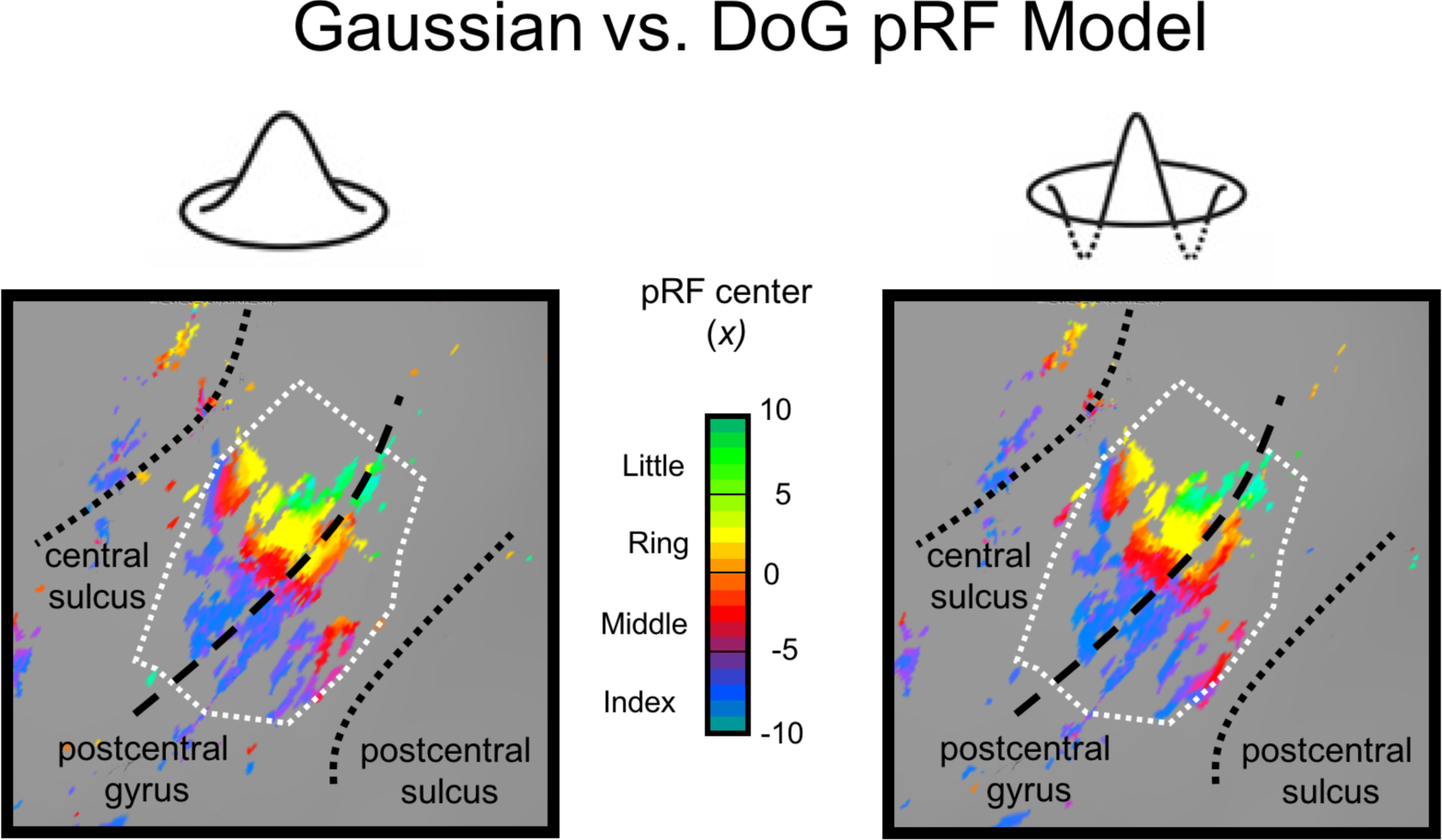
Gaussian vs. DoG pRF center maps for Subject 1.

